# Quotidian Profile of Vergence Angle in Ambulatory Subjects Monitored with Wearable Eye Tracking Glasses

**DOI:** 10.1101/2022.09.14.506830

**Authors:** Mikayla D. Dilbeck, Thomas N. Gentry, John R. Economides, Jonathan C. Horton

**Affiliations:** Program in Neuroscience, Department of Ophthalmology, University of California, San Francisco, San Francisco, CA 94143 USA

**Keywords:** pupil, ductions, strabismus, convergence insufficiency, interpupillary distance, Tobii Pro 3

## Abstract

**PURPOSE:** Wearable tracking glasses record eye movements and fixations as ambulatory subjects navigate their environment. We tested the performance of eye tracking glasses under laboratory and real world conditions, to characterize the vergence behavior of normal individuals engaged in their customary daily pursuits.

**METHODS:** To define the accuracy and variability of the eye tracking glasses, 4 subjects fixated with the head stabilized at a series of distances corresponding to vergence demands of: 0.25, 0.50, 1, 2, 4, 8, 16, and 32°. Then, 10 subjects wore the eye tracking glasses for prolonged periods while carrying out their normal activities. Vergence profiles were compiled for each subject and compared with interpupillary distance.

**RESULTS:** In the laboratory the eye tracking glasses were comparable in accuracy to remote video eye trackers, outputting a mean vergence value within 1° of demand at all angles except 32°. In ambulatory subjects the glasses were less accurate, due to tracking interruptions and measurement errors, only partly mitigated by application of data filters. Nonetheless, a useful record of vergence behavior was obtained in every subject. Vergence angle often had a bimodal distribution, reflecting a preponderance of activities at near (mobile phone, computer) or far (driving, walking). Vergence angle was highly correlated with interpupillary distance.

**CONCLUSIONS:** Wearable eye tracking glasses provide a history of vergence angle and the corresponding scene witnessed by ambulatory subjects. They offer insight into the diversity of human ocular motor behavior and may become useful for diagnosis of disorders that affect vergence, such as convergence insufficiency, Parkinson disease, and strabismus.

## INTRODUCTION

Vergence eye movements rotate the globes in opposite directions to align the foveae on visual targets located at a range of distances from the observer.^1^ They often incorporate changes in gaze direction, generating complex eye movements that combine saccades with shifts in vergence angle.^2^ Although such eye movements have been studied intensively in the laboratory, less is known about ocular motor behavior while humans navigate their visual environment in the course of normal daily life.

Many clinical disorders result from impairment of vergence function.^3^ In divergence insufficiency, subjects are able to fuse at near, but become esotropic at distance.^4–8^ In convergence insufficiency, fusion is intact at distance but a symptomatic exophoria (or even exotropia) emerges at near. This condition occurs in up to 4% of school children.^9, 10^ It is also common after head trauma or in certain neuro-degenerative conditions, such as Parkinson disease.^11–14^

To diagnose vergence disorders and their response to treatment, clinicians typically measure the near point of convergence and document the alignment of the eyes at near and far with a cover test.^15–17^ It would be valuable to obtain additional information, about the range, accuracy, and prevalence of vergence eye movements to targets located at difference distances. The advent of remote (e.g., desk mounted) video-based eye trackers has made it relatively easy to acquire such data in a laboratory setting.^18^ However, the data are usually collected over a relatively brief period and do not reflect vergence behavior while engaged in a repertoire of regular activities.

Eye tracking glasses that can be worn while head-mounted allow one to record gaze direction of ambulatory subjects moving through their visual environment, as captured by a scene camera. Here we described our experience with Tobii Pro Glasses 3 in the measurement of vergence eye movements. The device provides a readout of binocular gaze position, but information about the position of each eye alone is extractable, allowing one to calculate vergence angle.

This study consists of two halves. In the first part, we test the performance of the instrument in a series of laboratory experiments, to define its accuracy and operational characteristics under optimal recording conditions. In the second part, we present data from a cohort of 10 normal subjects, capturing their eye movements and shifts in vergence angle over many hours of ambulatory recording. Such data were not attainable prior to the invention of wearable eye trackers. They provide the first documentation of human vergence behavior measured during normal activity for long periods. The instrument may permit quantitative assessment of function in patients with disorders of vergence, while the individuals are engaged in the actual visual tasks that present a challenge to them.

## METHODS

### Vergence Angle Measurements

These experiments were conducted with the Tobii Pro Glasses 3 (www.tobiipro.com), a third generation instrument that consists of eye glasses with 8 infrared illuminators and 2 cameras embedded in each plano lens (Fig. 1). The device incorporates a scene camera that captures 106° horizontally by 63° vertically at 25 Hz, along with a microphone, accelerometer, gyroscope, and magnetometer. Data are streamed from the glasses via a cable to a recording unit worn by the subject. In the recording unit the position of each eye relative to the scene is computed at 50 Hz from information about the location of the pupil center and the 8 illuminator reflections on the cornea. The fixation point is overlaid atop the scene camera view and the composite image is viewable in real time via a wireless connection to a tablet, computer, or smartphone. Data are also serialized on a secure digital (SD) card in a variety of file formats: JSON, mp4, and csv.

**Fig. 1.** Composite of images taken under infrared and visible light of a subject wearing the Tobii Pro Glasses 3. The device contains 8 miniature infrared illuminators and two cameras (inferonasal) embedded in each lens. The 8 illuminators are reflected on the cornea in a semi-circular pattern. A camera mounted on the bridge of the glasses captures the scene. A microphone is located just above the camera.

The Tobii Pro Glasses are powered by lithium ion batteries with a lifetime of 100 min. An external rechargeable battery pack was plugged into the recording unit to allow up to 12 hours of operation. Both devices were placed into a lightweight satchel to allow the subject unrestricted mobility. When outdoors, subjects wore a pair of slip-on tinted infrared-blocking lenses. The Tobii Pro Glasses were calibrated by having subjects fixate a bull’s eye target held between 50-100 cm. Spherical lenses were inserted into the glasses to correct refractive error.

Using a custom script written in Igor Pro (www.wavemetrics.com), data were extracted from the JSON file encoding the three dimensional vector representing the direction of each eye. Horizontal eye positions in degrees were calculated from these vectors by applying the following functions:

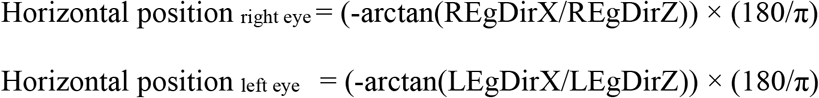

Interruptions and aberrant points in the data due to transient loss of eye tracking in one or both eyes were corrected by applying two filters.^19^ The first filter filled in gaps lasting up to 25 samples with the median of the surrounding 24 samples. The second filter removed spurious readings by comparing the value of each point with the 24 points surrounding it. If more than 1° outside the median, it was replaced with the median value.

To calculate vergence, the horizontal position of the right eye was subtracted from the horizontal position of the left eye. Positive values denoted convergence. Histograms were compiled in 0.2° bins showing the time that each subject fixated at a given vergence angle.

The JSON file also provides a gaze origin variable for each eye, measured at 50 Hz, which represents the distance of the pupil center from the cyclopean axis. Interpupillary distance was derived by adding the absolute values of each gaze origin variable. The distribution of interpupillary distances was determined over the duration of each recording.

### Experimental Subjects

This study was approved by the UCSF Institutional Review Board and followed the principles of the Declaration of Helsinki. Informed consent was obtained from adult subjects. Minors provided their assent and a parent gave informed consent.

In the first part of this study, the functional capability of the Tobii Pro Glasses 3 instrument to measure vergence angle was defined in laboratory experiments conducted in 4 adult subjects, several of them authors of this paper. In the second part, 10 normal subjects ranging in age from 10-67 years wore the eye tracking glasses for a prolonged period while going about their daily activities. All subjects had normal visual function, verified by ophthalmological examination, including assessment of acuity, pupils, eye movements, stereopsis, and fundi. Subjects with pathological nystagmus, strabismus, corneal disease, or prior ocular surgery were excluded. No refractive correction was necessary for the subjects testing the performance of the Tobii Pro Glasses in the laboratory, but some subjects participating in ambulatory monitoring wore soft contact lenses or spherical corrective lenses that fit into the glasses frames.

For the laboratory testing, each subject sat in a chair with the head immobile in an adjustable chin/forehead rest mounted on a table that could be moved vertically. The room was lit with fluorescent lights at a typical indoor brightness level (500 lux). The tracker was found to perform erratically in dim light, presumably because the dilated pupil is clipped by the upper eyelid. It also performed unreliably in direct sunlight, unless the infrared-blocking lenses were worn, because the corneal reflections from the illuminators were washed out by solar light.

Each subject fixated a crosshair target mounted on a tripod placed at the appropriate distance for a series of vergence angles: 0.25, 0.5, 1, 2, 4, 8, 16 and 32°. Subjects’ interpupillary distances were measured manually using a ruler (60, 64, 65, 68 mm) and also reported by the eye tracking glasses (60.5, 64.5, 64.1, 67.7 mm) during distance fixation. The value provided by the glasses was used to calculate the viewing distances for each subject’s series of vergence angles:

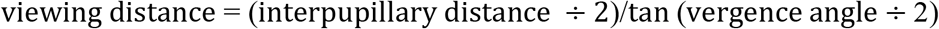

Specific information about each experiment is provided as the data are described in the Results.

For the ambulatory recordings of vergence angle, 10 subjects were asked to wear the eye tracking glasses for as long as they were willing, while engaged in their normal routine over the course of a day. Some subjects reported discomfort from the hard plastic nosepiece, due to the weight of the eye tracking glasses (60 g), sunglasses (30.7 g) and corrective lenses (17.3 g). The right temporal piece of the glasses became hot during prolonged recordings, bothering some subjects. Otherwise, the glasses did not interfere with routine activities, such as driving, running, shooting baskets, watching television, working at a computer, etc. Subjects were instructed to remove the eye tracking glasses before entering a lavatory. When placed back on the head, they immediately resumed tracking with the same calibration.

## RESULTS

**Figure 2A** shows a 15 min recording of a subject fixating a target at 14.68 m, corresponding to a vergence angle of 0.25°. There was a slow conjugate 4° rightward drift in eye position. This was unexpected, because the subjects in these experiments fixated the crosshair so faithfully that visual fading occurred for objects located in their peripheral visual fields. The most likely explanation is that the subject’s head turned slightly leftward during the recording. The mean vergence angle reported by the instrument was 3.0 ± 0.5°, representing an error of 2.75°. This means that although the subject was fixating at 14.68 m, the instrument recorded that he was fixating at 1.22 m. Splitting the recording into individual 1 min epochs showed a family of histograms, each narrower than the overall 15 min envelope (**Fig. 2 B,C)**.

**Fig. 2.**
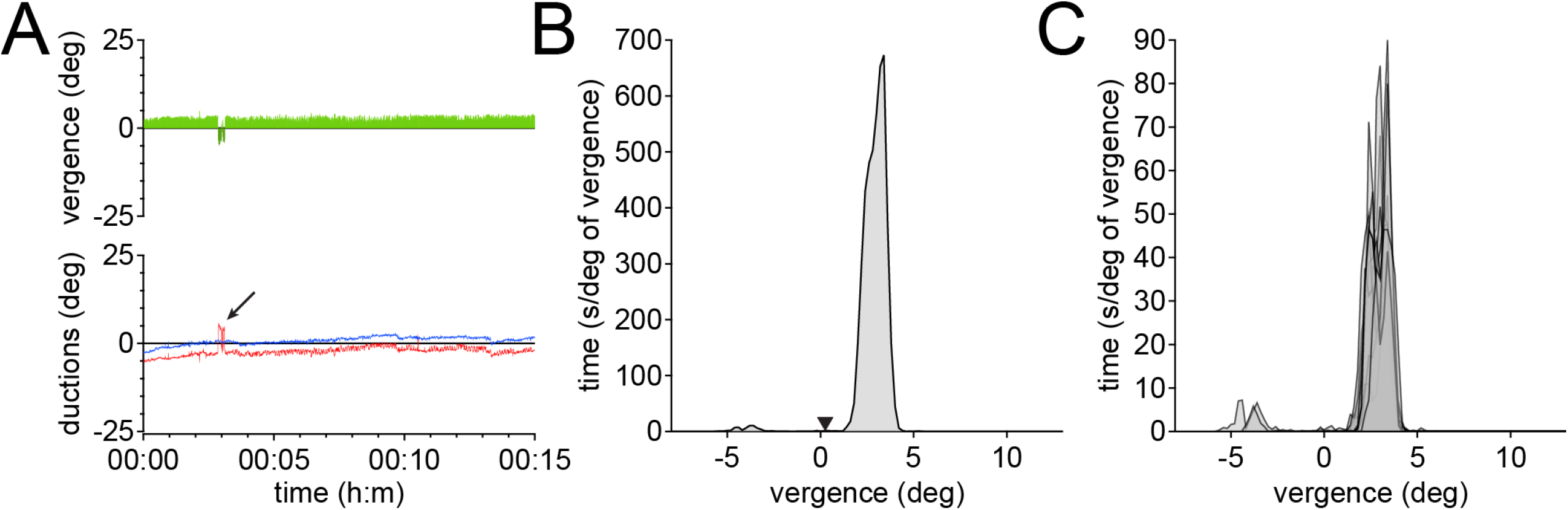
Vergence angle during steady fixation on a crosshair target at 14.68 meters for 15 minutes after calibration at a distance of 75 cm. **A)** Ductions (positive values denote right gaze) show a mostly parallel drift of about 4° in the position of each eye (red = right eye; blue = left eye). Coughing at 3 minutes (arrow) produced an artifact in the right eye trace, giving a transient negative vergence value. **B)** Histogram of vergence angle in (A), plotted in 0.2° bins (positive values denote convergence). The required vergence angle calculated from the subject’s interpupillary distance of 64.1 mm was 0.25° (arrow), but the mean value measured by the instrument was 3.0 ± 0.5° (range 1.3 - 4.6°). **C)** Same vergence data plotted in 15 sequential 1 minute intervals show slight fluctuations minute-by-minute in the measurements of vergence angle, with spurious readings at −5° from the cough.

**Figure 3A** shows a recording from the same subject, now converged at 16° on the crosshair. The right eye’s trace was more irregular than the left eye’s trace, with a 2 s interval marred by artifact. The mean vergence measurement was 15.7 ± 1.3°, a more accurate reading than recorded with the eyes diverged (**Fig. 3B**). Individual 1 min epochs showed jitter in the position of the peaks, as observed at 0.25° (**Fig. 3C**).

**Fig. 3.**
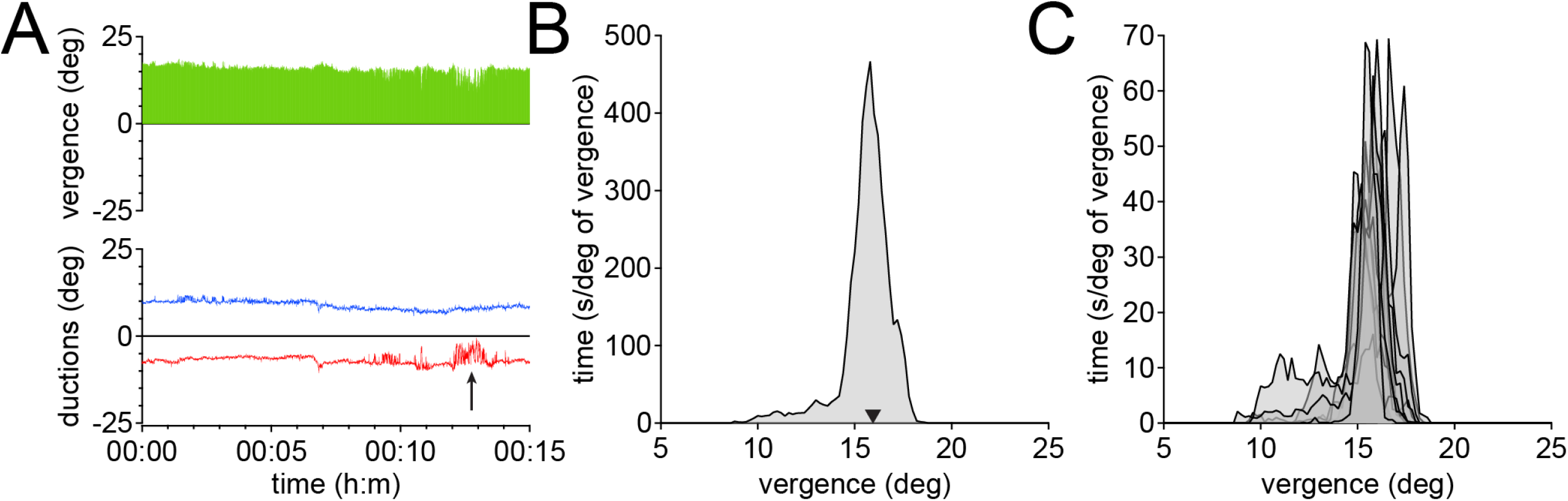
Vergence angle during steady fixation by the same subject in Fig. 2, viewing a crosshair at 22.8 cm for 15 min, requiring vergence of 16°. **A)** Ductions showed a slight outwards drift of the eyes over this interval, although the subject reported a single, fused fixation target throughout the recording. Artifact appeared in the right eye trace (arrow), without known cause. **B)** Vergence angle in (A) was 15.7 ± 1.3°. **C)** 15 sequential 1 minute plots of data, showing variability in measurements of vergence angle over time.

Given this difference in the accuracy of the eye tracking glasses at near versus far, we sought to determine if the distance at which calibration is performed is an important factor. The manufacturer recommends calibration of the eye tracking glasses while fixating between 50-100 cm. Testing we performed showed that calibration fails outside a range of 40-120 cm. **Figure 4A** shows a subject in whom calibration was obtained 5 times at 50, 75, and 100 cm. Subsequent fixation at distance after each calibration showed scatter in individual 1 min measurements of vergence angle. The scatter was unrelated to the calibration distance used to make the recording. **Figure 4B** shows a different subject, fixating at distance for 5 min epochs, after calibration performed at 75 cm. Individual peaks show scatter similar to that observed when calibration distance was varied. From these experiments we concluded that the choice of calibration distance between 50-100 cm does not matter. Furthermore, repeated determinations of vergence angle at distance in the same subject show a scatter in measured values that ranges over several degrees. For the sake of consistency, we decided to perform all subsequent calibrations at 75 cm.

**Fig. 4.**
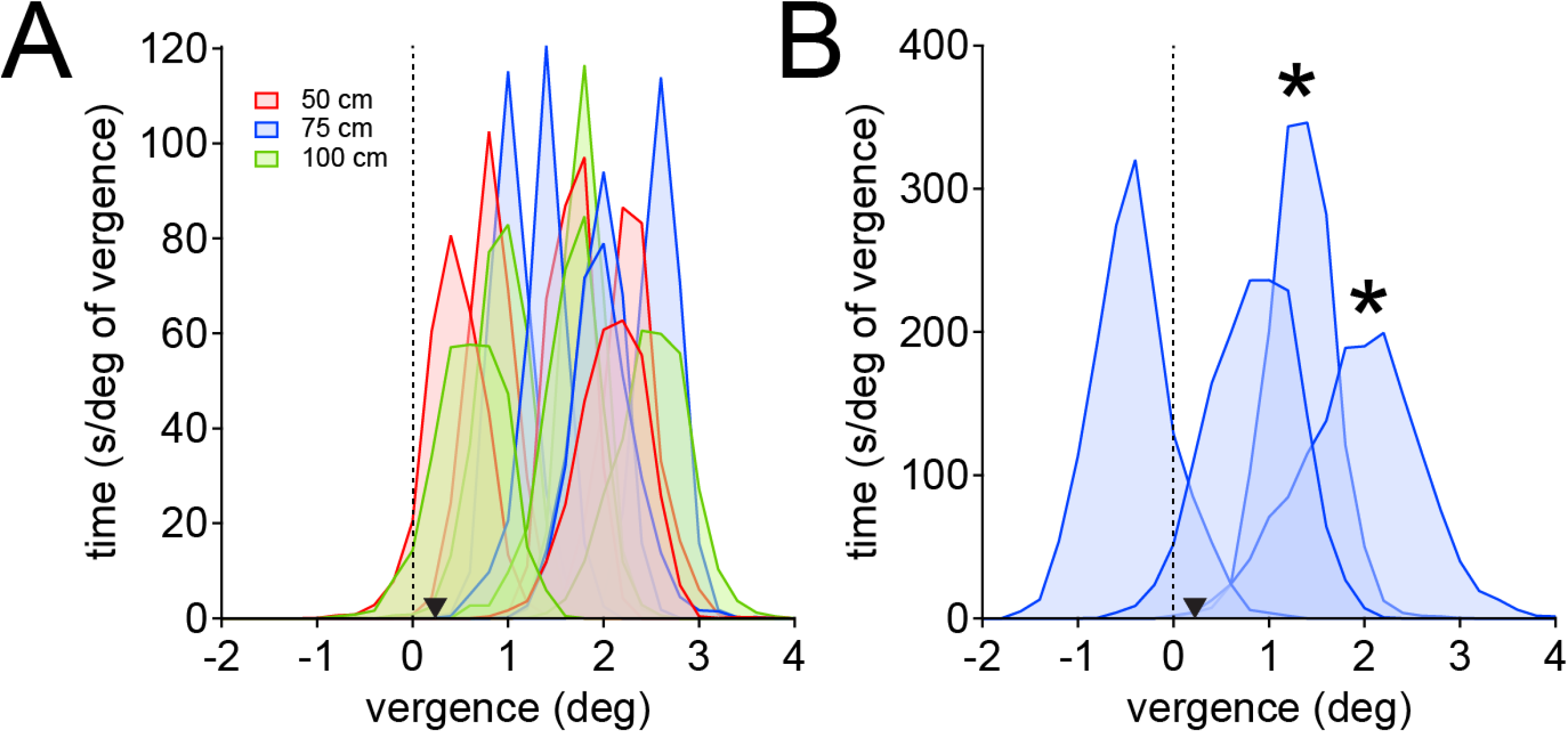
Calibration distance does not affect measurement of vergence angle by the eye tracking glasses. **A)** Histograms of vergence angle, recorded 5 times, for a 1 minute duration, each after a fresh calibration performed at either 50, 75, or 100 cm. The subject, with an interpupillary distance of 60.5 mm fixated at 13.87 meters, corresponding to a vergence demand of 0.25° (arrowhead). Scatter in measurements is unrelated to the calibration distance employed. The mean vergence was 1.7 ± 0.7°, an error of 1.45° **B)** Histograms of 4 epochs, each lasting 5 min, from a different subject fixating at 15.52 meters for a vergence demand of 0.25° (arrowhead). Calibration was obtained at 75 cm; two peaks marked with an asterisk were performed with the same calibration. The mean values ranged from −0.32 to 2.05°.

**Figure 5** shows data from 4 subjects fixating a crosshair for 5 min at distances that correspond to a series of increasing vergence angles: 0.25 - 32°. The traces of ocular ductions contain noise, especially at 32° (**Fig. 5A**). The location and width of peaks is variable among subjects. This variability is illustrated in **Figure 6**. At distance, the eye tracking glasses usually reported values greater than the true vergence angle. At 32°, they reported a mean value of only 28.9°. Subjects denied diplopia at 32°, even after 5 min of sustained convergence, so the discrepancy was likely due to tracker inaccuracy rather than inability to maintain a highly converged posture.

**Fig. 5.**
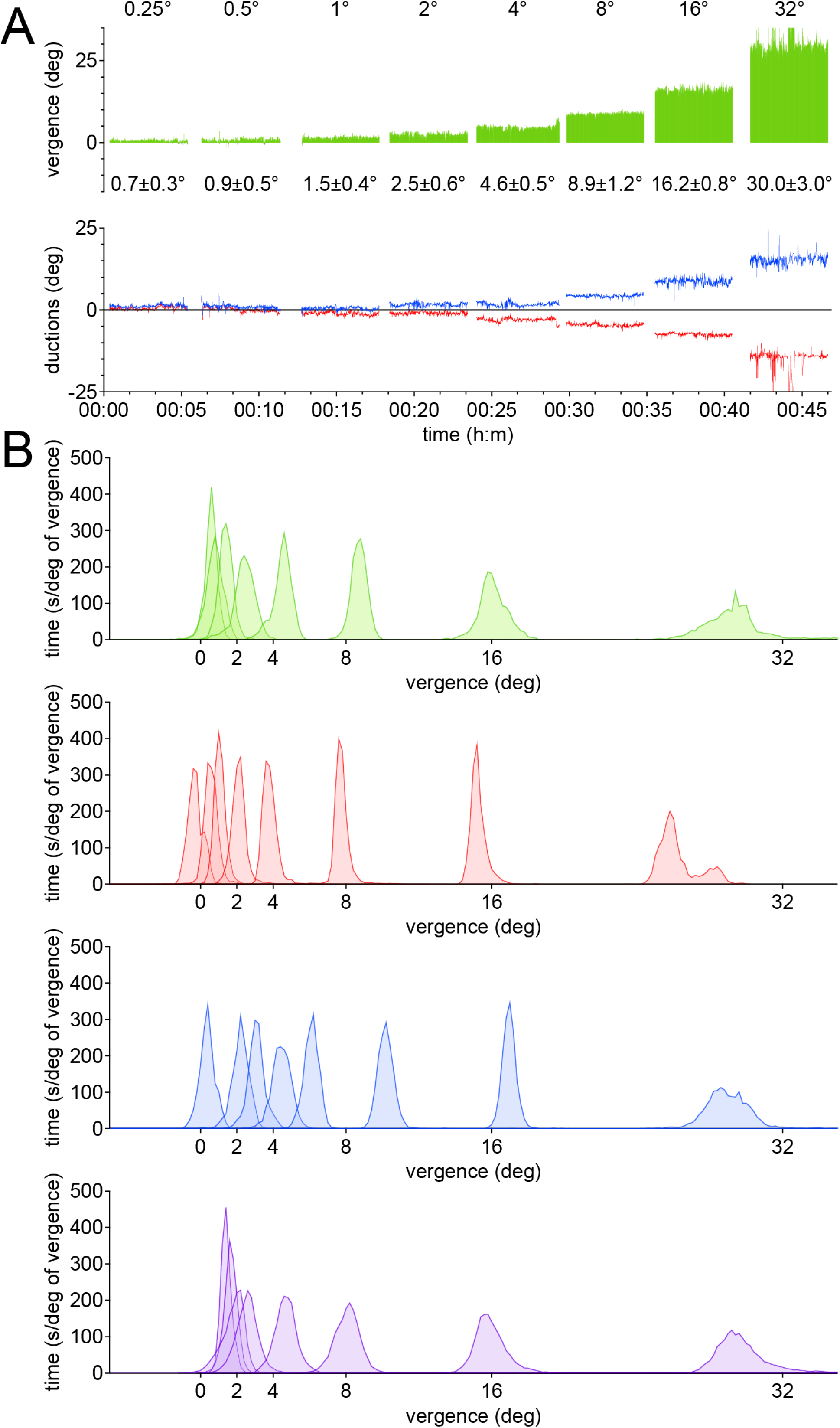
Variation in vergence angle measurements among subjects. **A)** Duction traces from a subject fixating at decreasing distances corresponding to vergence demand of 0.25, 0.5, 1, 2, 4, 8, 16 and 32° (top row). Each epoch lasted 5 minutes; gaps in the traces represent brief intervals between each fixation period. Tracker noise was most evident at 32° vergence angle. Mean vergence is listed below each plot. **B)** Data from 4 subjects (top series from subject in (A)), fixating for 5 minutes at each distance corresponding to the 8 vergence angles. Measurements deviate from the true vergence angle for some trials, and this error varies from subject to subject.

**Fig. 6.**
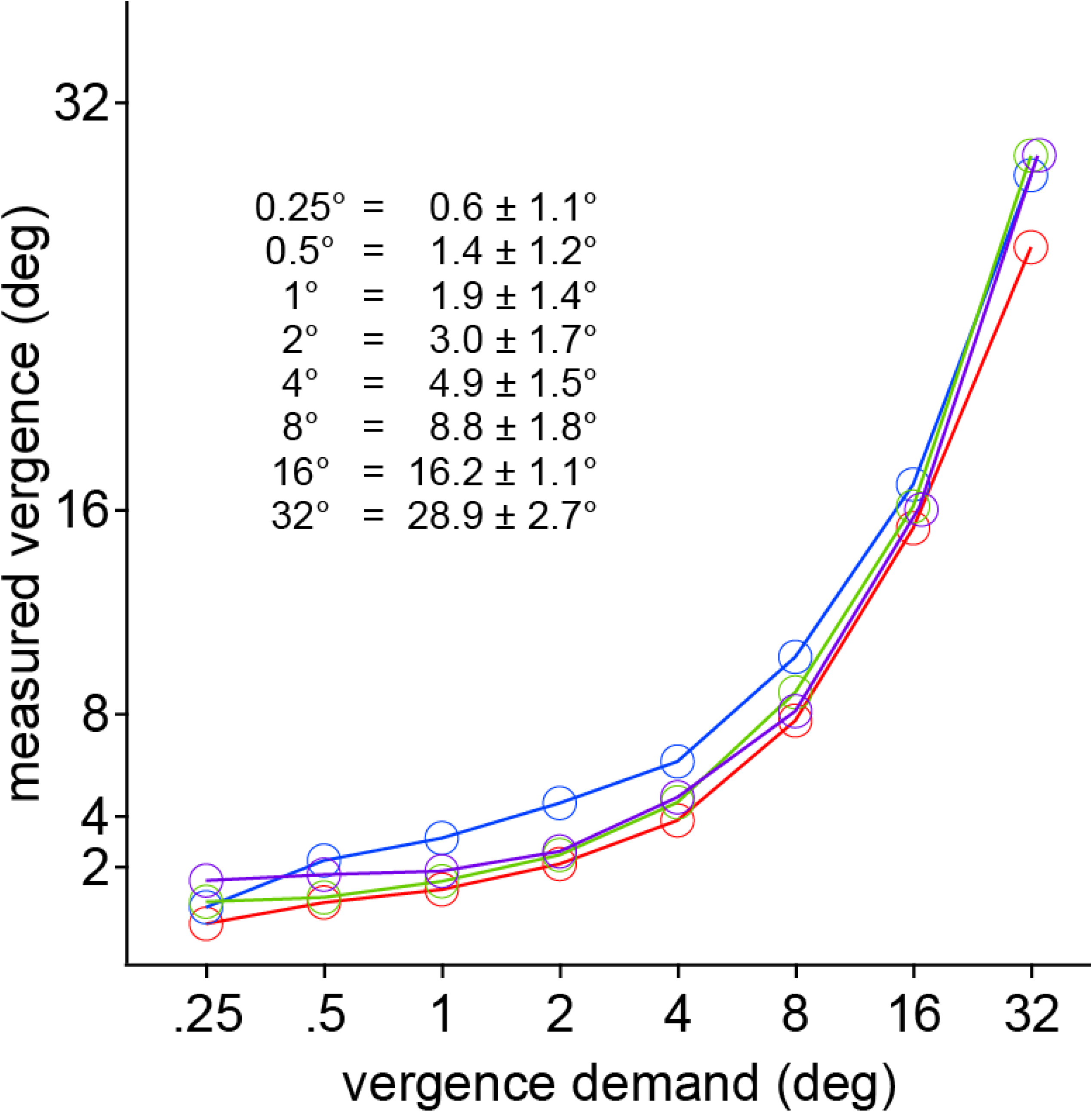
Plot of vergence demand versus measurement rendered by the eye tracking glasses. Data from Fig. 5, showing mean of the measured vergence at a range of vergence demands (0.25 - 32°) in each subject. Inset table shows (left) vergence demand and (right) measured mean vergence.

**Figure 7** tests the stability of the output provided by the eye tracking glasses over a prolonged time period. The subject fixated the crosshair at distance, converging at 0.25° for 60 s, every 15 min over 2 hours. Between measurements he worked freely at his computer. The mean vergence angle measured during the 9 distance fixation epochs was 2.0 ± 0.3°. Although there was an offset of 1.75° from the vergence demand, the data from each 60 s epoch showed no systematic drift over time (**Fig. 7C**).

**Fig. 7.**
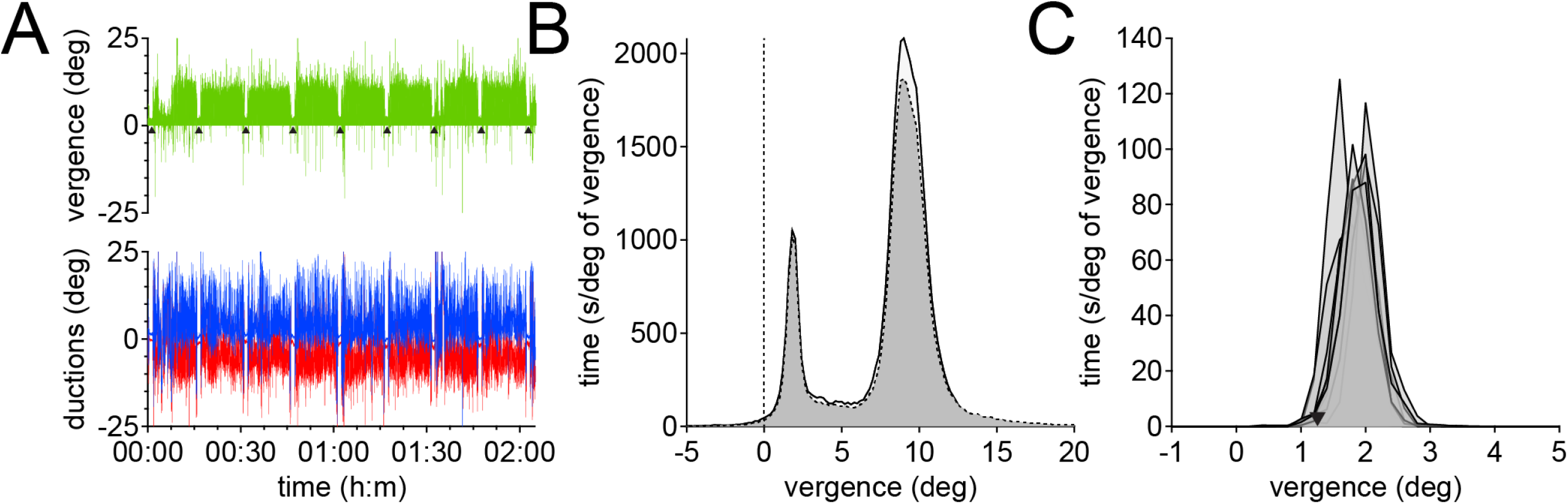
Stability of vergence angle measurement over time. **A)** The tracking glasses were worn by a subject (same individual illustrated in Fig. 4A) for 2 hours, while working at a computer. Every 15 minutes (arrowheads) he was asked to fixate for 60 s a target at 13.87 meters, vergence demand 0.25°. **B)** Histograms showing vergence angle over 2 hours. The bimodal distribution reflects epochs working at near on the computer and fixating at distance on a target with the head stabilized. Dashed line shows unfiltered data; solid line shows data after application of median filters. **C)** Plot of 9 histograms, each containing 60 s of data while converged at 0.25°, showing the consistency of readings compiled over 2 hours. The mean vergence measured 2.0 ± 0.3°, corresponding to an error of 1.75°.

**Figure 8** explores the impact of the median filters on the raw data traces. In this 5 s data excerpt, noise is present in the instrument’s position signals, even though the recording was made under optimal, head-stabilized conditions. As a result, the eye tracking glasses sometimes falsely reported a divergent (exotropic) alignment during fixation at distance. Short gaps occurred frequently in the position signal from one or both eyes, which were filled in by the first median filter. The second median filter is intended to mitigate inaccurate position signals, which often surround gaps in the recording traces. However, application of the filter sometimes exacerbated recording artifacts (compare **Fig. 8A** and **B**), especially when unequal gaps occurred in the record of each eye, in conjunction with an abrupt change recorded in the position of only one eye.

**Fig. 8.**
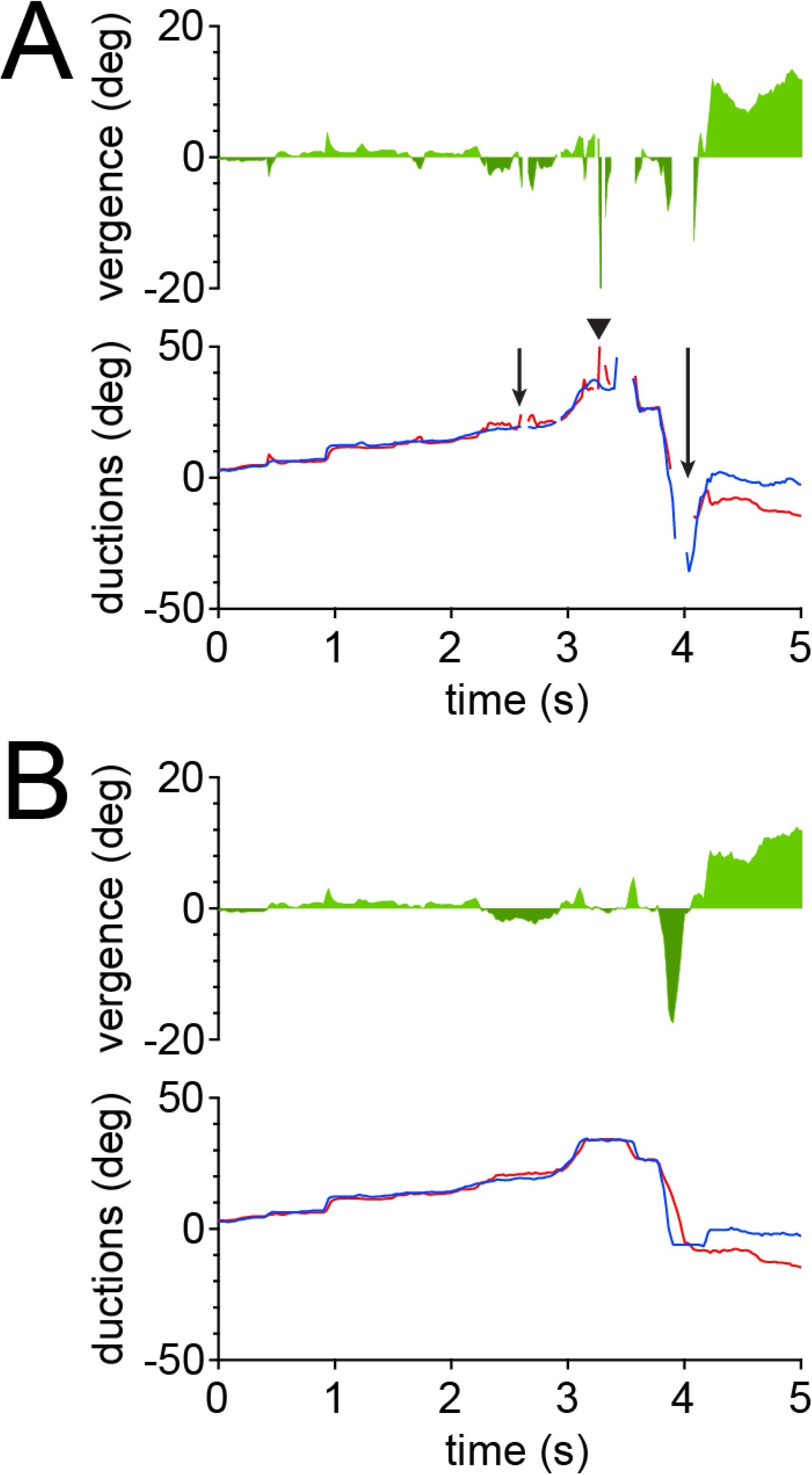
Effects of the median filters. **A)** 5 s of data from Fig. 7A starting at 1:33:05 showing impact of applying the median filters. At 2.3 s, inaccurate right eye position signals from the eye tracking glasses produce a spurious exotropia (short arrow). At 3.3 s, inaccurate right eye position signals recur (arrowhead), causing another false exotropia reading, a spike of −20°. Later, several breaks occur in both traces (long arrow), producing gaps in the vergence angle reading. **B)** After application of the median filters, the data traces become continuous and the spurious exotropia spike of −20° is eliminated. Filling in subsequent gaps in the traces, however, exacerbates a large, artifactual exotropia deflection. This occurs because the slope of the right eye’s trace, relative to that of the left eye, is altered by application of the median filters.

**Figure 9** shows data from a subject recorded in the second portion of this study, under free ranging conditions. After tracker calibration at 75 cm, the subject fixated for 1 min at a series of distances that corresponded to vergence angles from 0.25 - 32°. The distances were measured with a retractable steel tape measure and the fixation target was handheld. The purpose of this procedure was to obtain recordings at known, fixed vergence demand to correct any errors in the vergence angle reported by the eye tracking glasses. Under these field conditions, measurements (**Fig. 9A**) were less accurate and more variable than those obtained in the laboratory (**Fig. 5A**).

**Fig. 9).**
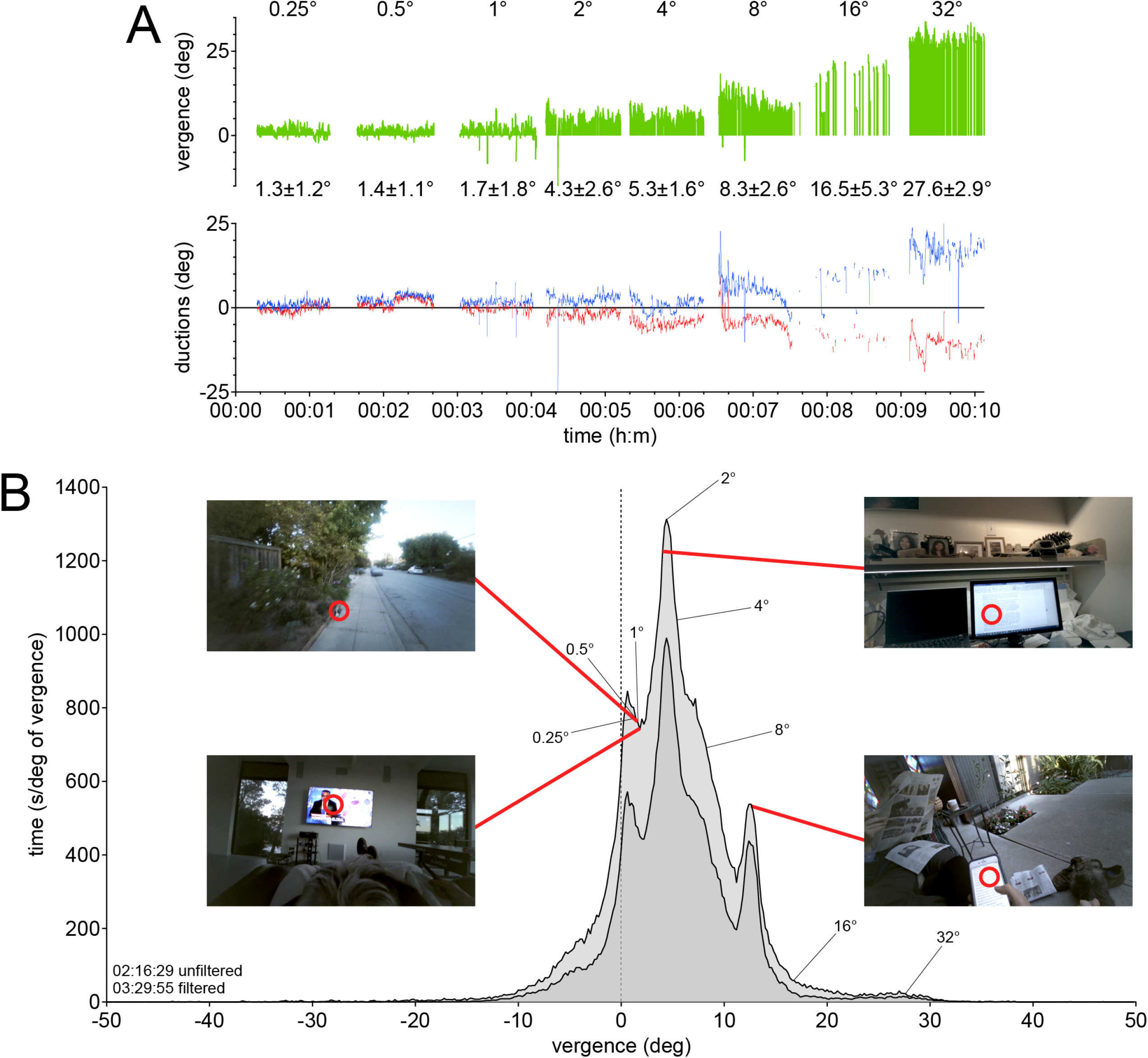
Vergence angle profile of a 52-year-old ambulatory subject. **A)** Before the recording, her interpupillary distance was measured manually and then distances were calculated for target presentation at vergence angles ranging from 0.25-32°. The traces show more gaps, drift, and spurious points than data gathered in the lab (Fig 5A). Numbers in top row denote vergence demand; numbers underneath indicate measured mean vergence. **B)** Histogram showing amount of time spent at each vergence angle. Dark shading represents unfiltered data. Mean value of data measured at each vergence demand in (A) is shown by thin black lines. Red lines point to mean value of vergence angle during the activity shown in the scene camera frame.

Over a total recording time of 4:14 hours of the subject engaged in various activities, reflected by the profile of her vergence behavior (**Fig. 9B**). The unfiltered data had a duration of 2:16:29, which increased to 3:29:55 after application of the median filters. The filtered data were shorter than the total recording time because the subject took rest breaks and there were interruptions in the eye position signals that lasted more than 25 samples. Application of the filters “rescued” a greater percentage of the recording time during field recordings than during laboratory recordings (**Fig. 7B**). This difference reflected that fact that cleaner data were obtained in the laboratory, so application of the filters had less impact. The filters greatly increased the duration of usable field recordings, without changing the overall shape or location of peaks in any subject’s vergence profile.

Correlation with the video from the scene camera revealed that some of the distinct peaks were generated mostly by a single activity, such as viewing a smartphone or a computer monitor. Other activities, such as walking a dog or viewing a television, occurred at small vergence angles. They did not correspond uniquely to a single peak, because a mixture of many different behaviors happened at a small vergence angle. Another factor is that vergence angle changes less than 2° between infinity and 2 meters. Given the limits of the tracker’s accuracy, various activities conducted at slightly different distances within this range were merged in the subject’s vergence profile.

Negative vergence values present in the subject’s vergence profile (**Fig. 9B)** signify exotropia. Clinical examination revealed that this subject was orthotropic at distance. Therefore, all points graphed to the left of 0° represented inaccurate measurements of her vergence angle. Artifactual negative values occurred while subjects were fixating at distance (**Fig. 8**) and also at near (**Fig. 7**).

**Figure 10** shows a family of vergence angle distributions compiled from the other 9 subjects during prolonged ambulatory recordings. All the subjects were orthotropic at distance fixation. Therefore, as in **Figure 9**, all negative vergence values were a product of instrument error. The individual profiles varied considerably, because each person was engaged in different pursuits during the recording session. In general, there was a tendency for subjects’ vergence behavior to exhibit two modes. Subjects fixated predominately at near or at far, with a relative paucity of intermediate points.

**Fig. 10).**
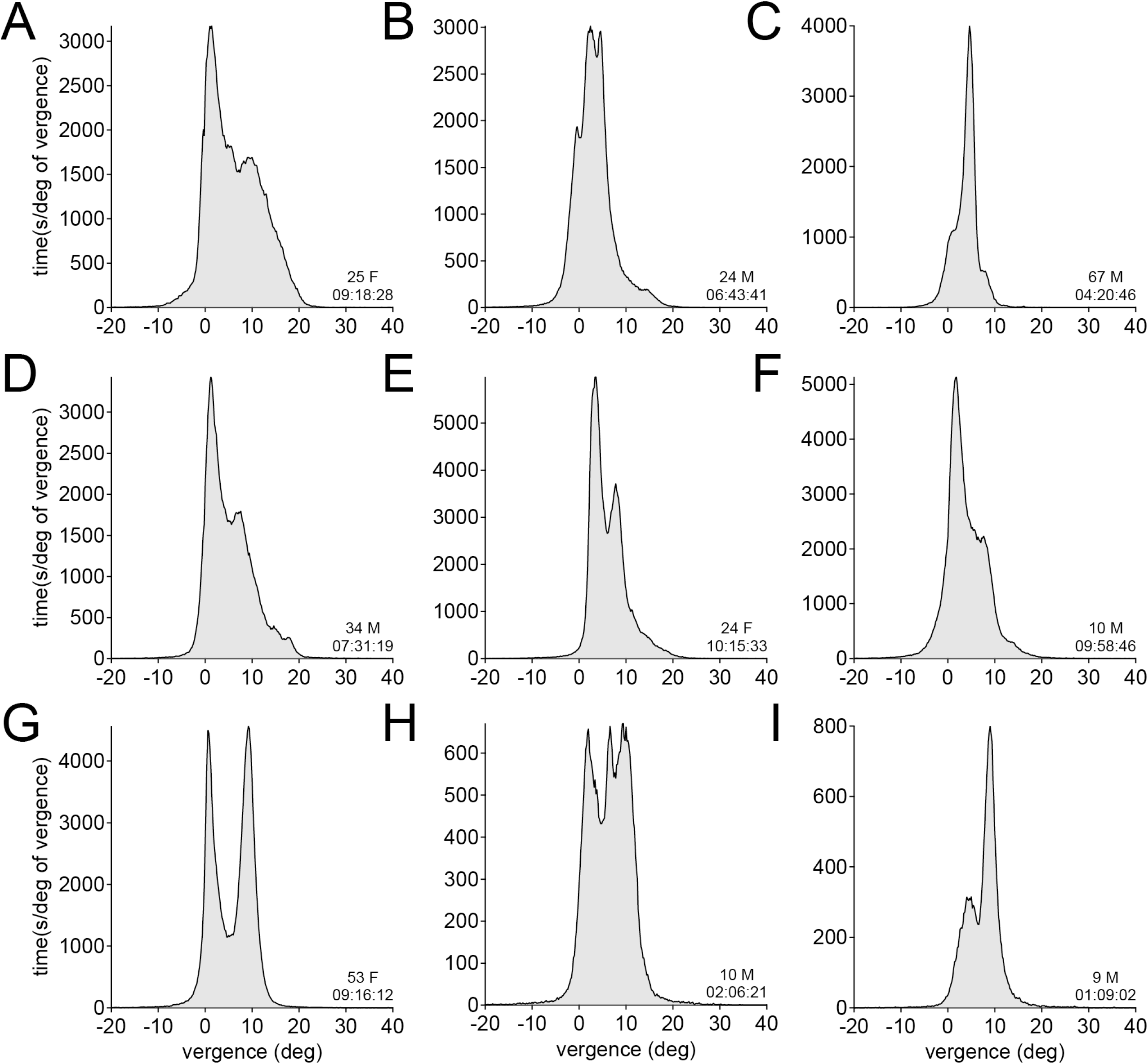
Variability of vergence angle profiles. **A - I)** Vergence angle histograms from 9 different subjects. Age, gender, and duration of filtered data are listed. Each profile is unique to the repertoire of activities in which the subject was engaged during the recording.

**Figure 11** shows plots of the 9 subjects’ interpupillary distances measured during the same ambulatory recording sessions. In each subject, the interpupillary distance ranged over about 4 mm. The plots resembled the vergence profiles (**Fig. 10**), which was expected, because the tracking glasses incorporate data about the location of the pupil center in their calculation of eye position.

**Fig. 11).**
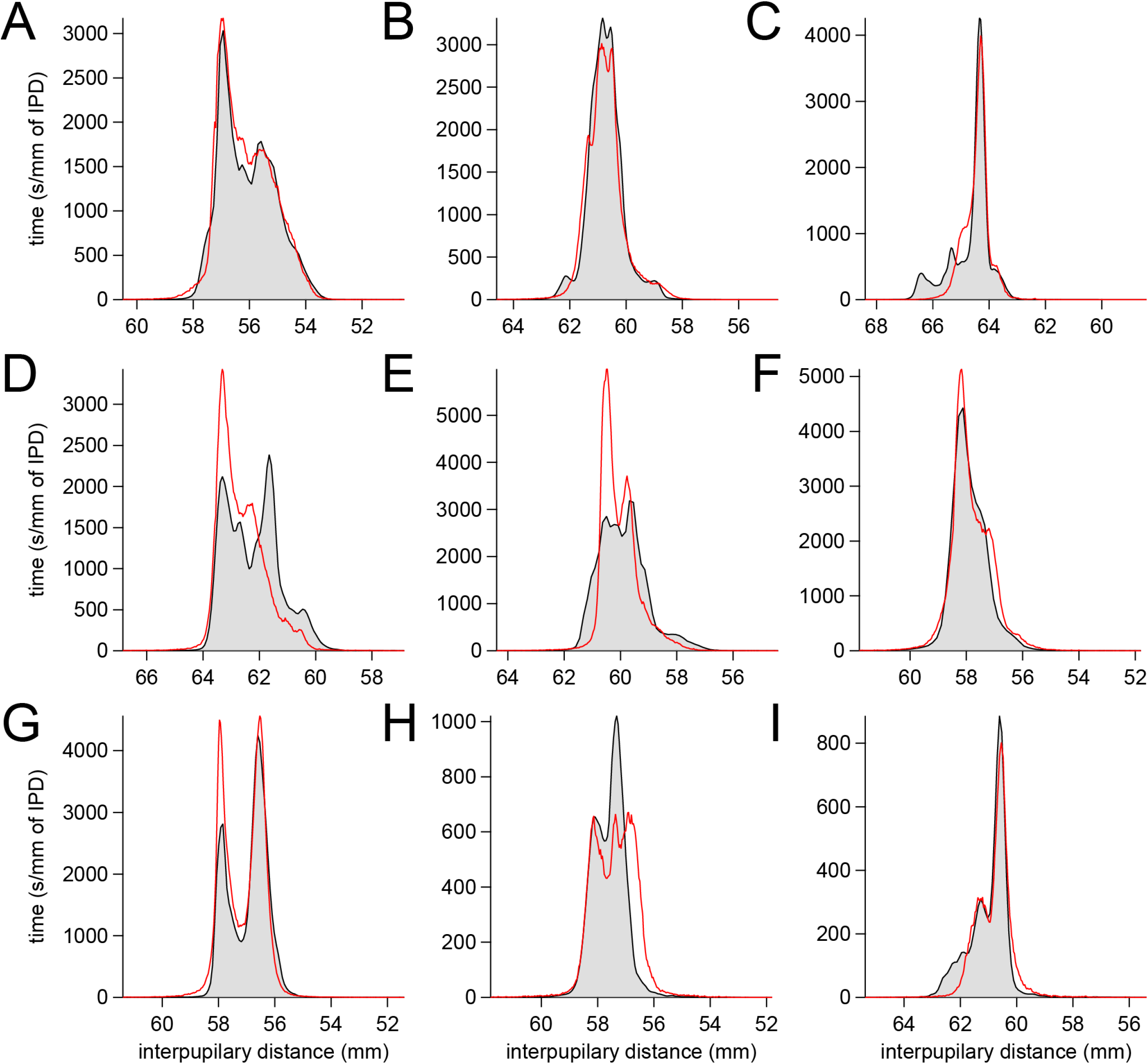
Interpupillary distance histograms for 9 ambulatory subjects. **A-I)** Same subjects shown in Fig. 10, plotting the interpupillary distances at each moment during their recording session. Highest values corresponds to the interpupillary distance in primary gaze, fixating at distance. Interpupillary distance ranged over 4 mm in most subjects and usually resembled the profile of vergence angle (red outlines, from Fig. 10).

**Figure 12** illustrates the relationship between vergence angle and interpupillary distance for Subject A, sampled at 50 Hz over the course of more than 9 hrs. **(Fig. 10, 11)**. There was a negative correlation (r = −0.82), with each 1 mm decrement in interpupillary distance corresponding to a 5.1° increase in vergence angle. For the 9 subjects, r = −0.78 ± 0.06 and the mean increase in vergence angle was 4.9 ± 1.4° per 1 mm decrease in interpupillary diameter.

**Fig. 12).**
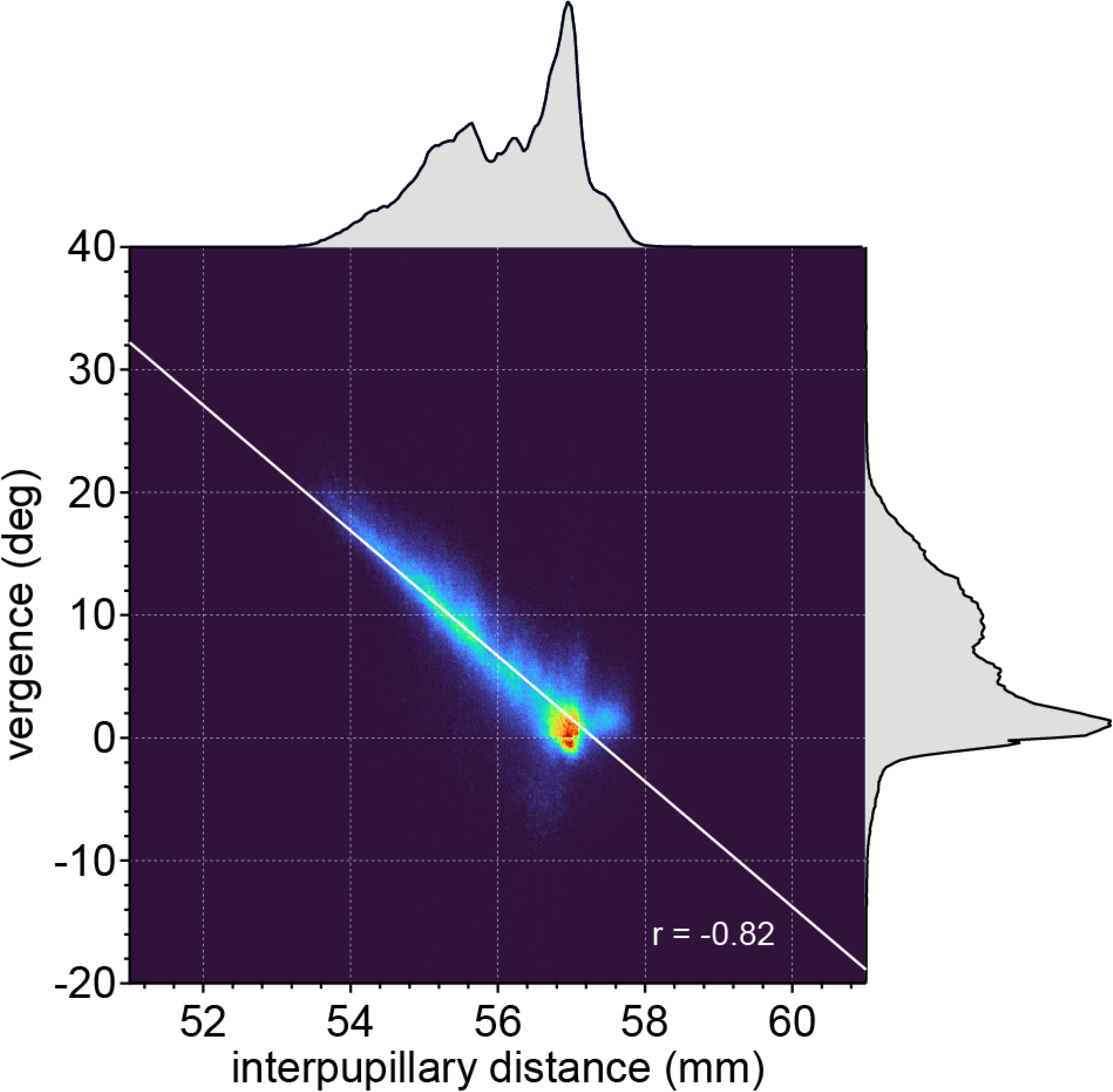
Correlation between vergence angle and interpupillary distance. Data from subject A in Fig. 11, plotting interpupillary distance (~ 1.7 × 10^6^ samples) versus the simultaneously measured vergence angle. The Pearson correlation coefficient was −0.82. The histogram of the interpupillary distance is shown above; the histogram of the vergence angle is shown at right.

## DISCUSSION

The advent of eye trackers that can be worn by ambulatory subjects while engaged in their normal activities has opened a new field of investigation into human behavior. They permit one to compile data about eye movements and fixations made by individuals as they explore their environment over long periods in unconstrained settings. In this report we show data from 10 normal individuals, capturing a history of the angle at which their eyes were converged (**Figs. 9, 10**). Each profile was different, idiosyncratic to the tasks and actions of the subject, but several features were consistent. A large fraction of time was spent fixating at near targets, such as a mobile phone or computer monitor (or even a book). Much of the remaining time, fixation occurred between infinity and several meters. Fixation was relatively less frequent in the intermediate range, from just beyond the outstretched hand to several meters away. As a result, for many individuals the profile of fixation distances had a bimodal distribution.

Ambulatory eye trackers have been useful for evaluation of performance in work settings. For example, they have been deployed to monitor the gaze of nurses in an intensive care unit, chefs cooking food, and workers in industrial occupations.^20–22^ They may also prove valuable for early detection of impaired eye movements associated with certain movement disorders, such as Parkinson disease and supranuclear palsy. In Parkinson disease, deficient convergence with diplopia at near is a prevalent symptom.^13, 23, 24^ In patients with autism and schizophrenia, abnormal patterns of eye movement behavior have been identified.^25–30^ It would be worthwhile to verify such observations, made in artificial laboratory settings, with recordings made in subjects’ natural environments. Ambulatory recordings of eye movements may also serve to diagnose and monitor patients with ocular misalignment or strabismus, especially when findings are present only intermittently.

It is important to evaluate the accuracy of face-mounted eye trackers to interpret the data obtained from ambulatory subjects. Accuracy refers to the offset between the fixation position measured by the tracker and the subject’s true fixation position.^31^ Remote video eye trackers tested on stabilized subjects in the laboratory have an accuracy between 0.5 - 1°, measured in the X,Y plane.^32–38^ Vergence angle is a measurement in the Z axis, but is derived simply by subtracting X axis (duction) variables. Our measurements of the face-mounted Tobii Pro 3 eye tracking glasses, also tested under optimal conditions, yielded a comparable accuracy of 0.5 - 1° (**Fig. 6**), except at the highest vergence angle (32°). Usually, the vergence angle reported was slightly too high during distance fixation and too low at the closest fixation point.

When deployed in ambulatory subjects, the Tobii Pro 3 eye tracking glasses were less accurate.^39^ The calibration traces in a typical subject (**Fig. 9A**) showed an error of 4.4° at 32° convergence, and an error up to 2.3° over other tested angles. The traces were also much noisier, with fluctuation and dropout of the eye position signal caused by tracker error. Our original intention was to use the measurements of vergence angle at known fixation distances made prior to each ambulatory recording to correct offsets in the value of vergence angle subsequently reported by the tracker. However, this proved of limited value, for several reasons. First, the error between vergence demand and recorded vergence varied at each distance, so a non-linear data transformation would have been necessary. Second, it was difficult to measure fixation distances in the field recordings with sufficient precision.

In addition to accuracy, the variability of tracker output during steady fixation is an important index of instrument performance. In the laboratory, vergence angles of 0.25, 0.50, 1, 2, 4, 8, and 16° had a mean standard deviation ranging between 1.1 – 2.7° (**Fig. 6**), and an absolute variation in eye position between 2-4°. For the ambulatory recordings, there was no way to assess variability in tracker recording of vergence, because vergence angle was not controlled. One gains some sense of the device’s performance by observing the spread of reported vergence angles into the negative, non-physiological range. All negative values of vergence angle, signifying divergence of the ocular axis, were artifactual. Rather than displaying a sharp vertical cut off at 0° vergence angle, each subject’s trace showed a tapered spread of errant negative values (**Fig. 10**). Such false points were generated during epochs of near fixation and distant fixation (**Fig. 7**). The variability of tracker measurements reduced the fidelity of the data, but there is no suggestion that it altered the position of peaks in any subject’s profile.

The close inverse correlation (r = −0.78) between interpupillary distance and vergence angle (**Fig. 11**) arises because the pupil centers are anterior to the centers of rotation of the globes. With each 1 mm decrease in interpupillary diameter there was an increase in vergence angle of 5°. If one’s goal in recording subjects is simply to track vergence angle, rather than X,Y gaze position, it may be possible to employ a lower-cost device that relies only on detection of the pupil and measurement of its center position. However, the additional information provided by the corneal reflection of the illuminators allows registration of the eyes’ fixation positions with the view provided by the scene camera.

## Acknowledgments

This work was supported by grants EY029703 (J.C.H.) and EY02162 (Vision Core Grant) from the National Eye Institute and by an unrestricted grant from Research to Prevent Blindness. Jessica Wong assisted with computer programming. The authors have no conflict of interests.

